# *Culex (Culex) acharistus,* Root, 1927 (Diptera: Culicidae), a new record for Bolivia

**DOI:** 10.64898/2025.12.09.692992

**Authors:** Frédéric Lardeux, Philippe Boussès, Carolina Quezada, Audric Berger, Rosenka Lardeux, Claudia Aliaga, Libia Torrez

**Affiliations:** Instituto de Investigaciones Biomédicas e Investigación Social (IIBISMED), Universidad Mayor de San Simón, Facultad de Medicina, Calle Venezuela, Cochabamba, Bolivia; Unité Mixte de recherche MIVEGEC (Maladies infectieuses et vecteurs : écologie, génétique, évolution et contrôle), Institut de recherche pour le développement (IRD) - Université de Montpellier - Centre National de la recherche Scientifique (CNRS), 911 avenue Agropolis, Montpellier, France; Laboratorio de Entomología Medica (LEMUMSS), Universidad Mayor de San Simón, Facultad de Ciencias y Tecnología, calle Sucre y Parque La Torre, Cochabamba; Bolivia; Centre de coopération internationale en recherche agronomique pour le développement (CIRAD), Unité Mixte de Recherche INTERTRYP (Interactions Hôtes-Vecteurs-Parasites-Environnement dans les Maladies Tropicales Négligées dues aux Trypanosomatidés), CIRAD - IRD - Université de Montpellier, Campus international de Baillarguet - TA A-17 / G, 34398 Montpellier Cedex 5, France; Laboratorio de Entomología y Parasitología, Instituto Nacional de Laboratorios de Salud “Dr. Néstor Morales Villazón” (INLASA), Pasaje Rafael Zubieta 1889, La Paz, Bolivia

**Keywords:** *Culex acharistus*, Culicidae, South America, Bolivia, taxonomy, morphology, COI sequences

## Abstract

This study reports the first confirmed records of *Culex (Culex) acharistus,* Root, 1927 in Bolivia, based on both morphological examination and biomolecular identification using the cytochrome c oxidase subunit I (COI) marker. Specimens were collected in four localities representing diverse environmental settings: La Paz (urban environment, ≍3600 m), the nearby high-altitude city of El Alto (≍4000 m), Cochabamba (semi-rural and urban habitats, ≍2600 m), and Licoma a large village in the Yungas region (subtropical environment, ≍1900 m). The larval habitats where the species was found closely match those reported in neighboring countries where *Cx. acharistus* also occurs, ranging from typical small peridomestic sites to ponds, ditches, and river seepages in semi-rural areas. All identification methods yielded unambiguous results, confirming the species’ presence in Bolivia and extending its known distribution in South America. This finding highlights the importance of integrating morphological and molecular approaches to achieve accurate and reliable mosquito species identification.

## Introduction

Bolivia, located at the heart of South America, spans a broad range of altitudes and climates—from the high Andes to lowland Amazonian plains—supporting a variety of ecosystems. With a recognition of 17 distinct ecoregions (Olson *et al*. 2001) and up to 23 sub-ecoregions (Ibisch & Mérida 2003), Bolivia is considered a megadiverse country (Ibisch 2001), reflecting its exceptional biological richness. This environmental heterogeneity contributes to the country’s remarkable insect diversity, with over 13 700 species already documented and the total number expected to double as new taxa are discovered and described (Moraes & Sarmiento 2019). Among them, the mosquito family Culicidae remains understudied, despite its medical importance in the region. Only a few dated general surveys of Bolivian Culicidae are available (Pinto 1932; Cerqueira 1943; Shannon & Cerqueira 1943; del Ponte *et al*. 1951; Martínez & Prosen 1953; Martínez *et al*. 1960; Prosen *et al*. 1962/63) which initially contributed to the compilation of a list of about 156 species. The national Culicidae list has progressively expanded through successive records and species descriptions (Sirivanakarn 1979; Peyton E.L. *et al*. 1983; Peyton E. L. *et al*. 1984; Harbach *et al*. 1984; Pecor *et al*. 1992; Brelsfoard 2005; Brelsfoard *et al*. 2006; Lardeux Frédéric *et al*. 2009; Juri *et al*. 2022; Lardeux F. *et al*. 2024b), and as a result of these cumulative contributions, the current national checklist of Culicidae, compiled by one of the authors (PB), now comprises approximately 240 species and continues to evolve, including the species documented in this study (VECTOBOL 2025).

This article formally reports the presence of *Culex (Culex) acharistus,* Root, 1927 in Bolivia, based on combined morphological and molecular evidence.

## Material and methods

### Study area

*Culex acharistus* specimens were collected in three different areas of Bolivia: (1) the Cochabamba city area in the center of the country, (2) the La Paz and El Alto city area in the Altiplano region, and (3) in Licoma, a large village in the Yungas region (FIG. 1).

**FIG. 1.**
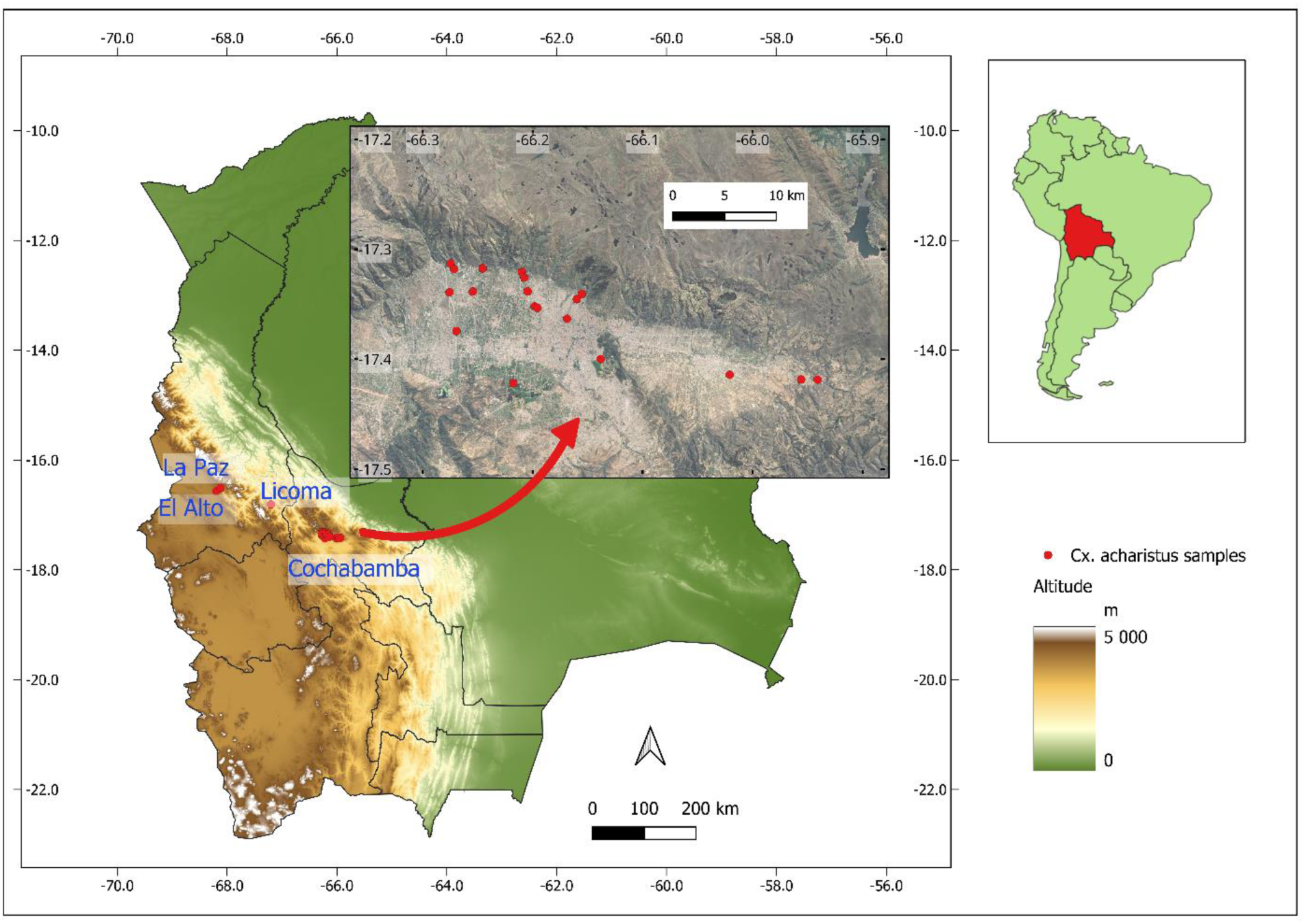
Sampling locations of *Cx. acharistus* in Bolivia. The main map shows the country, with two inset maps: the first highlights Bolivia within South America, and the second, overlapping Bolivia’s boundaries, shows the area of the city of Cochabamba.

The major city of Cochabamba (17°22’48” S, 66°09’36” W), with a population of around 800 000 (excluding the broader metropolitan area), is located in central Bolivia on a vast plain at an altitude of ≍2 600 meters above sea level. The city has a temperate climate with a mean annual temperature of about 18°C. Temperatures range from a cooler 10°C in the winter months of June and July to a warmer 26°C during the summer months of December and January. Precipitation is moderate, with the rainy season extending from November to March, yielding an average annual rainfall of approximately 700 mm. The collection sites were located either within the city of Cochabamba or in its surrounding areas, which include transitional zones with urban characteristics, as well as more rural environments featuring mixed housing, small-scale agriculture, and livestock settings, where both artificial and natural mosquito larval habitats often coexist.

In contrast, the La Paz/El Alto region, located at an average altitude of ≍3600 meters above sea level, has a more pronounced temperature variation and a cooler climate. La Paz (16°29’32”S, 68°07’15”W) is the administrative capital of the country, and with the contiguous city of El Alto (16°29’18”S, 68°12’19”W), the region totalizes about 2 million inhabitants. In La Paz, the average annual temperature is around 10°C, with winter temperatures dropping to as low as 0°C in June and July, while summer temperatures can reach up to 18°C in December and January. La Paz receives less precipitation than Cochabamba, with an average annual rainfall of about 550 mm, concentrated mainly between November and March. El Alto, at an elevation of ≍ 4000 meters has a colder climate with an average annual temperature of approximately 7.2°C. The warmest month is typically October, with average daytime temperatures around 14°C and nighttime lows near 5.4°C. El Alto experiences higher annual precipitation, totaling approximately 782 mm. Similar to La Paz, the rainy season spans from September to April, with January being the wettest month. The dry season from May to August sees significantly less rainfall, with June being the driest month. Both collecting sites were located in highly urbanized environments: one in the center of La Paz city, and the other in the El Kenko suburb of El Alto city.

Licoma (16°48’16”S, 67°12’11”W), a large village of about 1000 inhabitants, is located in the Inquisivi Province within the La Paz Department of western Bolivia. It is situated in the Yungas region, a transitional zone between the Andean highlands and the Amazonian lowlands, at an elevation of ≍1 900 meters. The climate is subtropical mesothermal, characterized by moderate temperatures throughout the year and distinct seasonality in rainfall. Mean annual temperatures range from 16°C to 22°C, with variations influenced by local microhabitats and slope orientation. The area receives between 1000 and 2000 mm of rainfall annually, primarily concentrated during the austral summer months (November to March), while the winter months (May to August) experience a pronounced dry season. The region is dominated by cloud forests and montane rainforests.

### Field mosquito collections

*Culex acharistus* was first collected in Bolivia in 2018 by one of the authors, when a specimen was fortuitously recovered from a bucket in a peridomestic setting in the Tiquipaya suburb of Cochabamba. However, this finding was not formally reported at the time. In the present study, we document this initial observation along with subsequent records of the species obtained during entomological surveys conducted in Cochabamba in 2020, 2023, and 2024, aimed at updating the mosquito species inventory for the area (Torrez *et al*. 2022; Quezada 2024). These surveys were part of the VECTOBOL initiative (Lardeux F. *et al*. 2024a), which is dedicated to improving knowledge of the geographic distribution of disease-vector insects across Bolivia. These surveys were carried out during both the rainy and dry seasons. In addition, *Cx. acharistus* larvae were incidentally collected in 2024 at two highly urbanized sites in peridomestic environment: the city center of La Paz and the El Kenko suburb of El Alto city. In the locality of Licoma, the species was identified from larval samples collected during a mosquito survey conducted by the Servicio Departamental de Salud (SEDES-La Paz) in the municipal cemetery (Table 1).

**Table 1.**
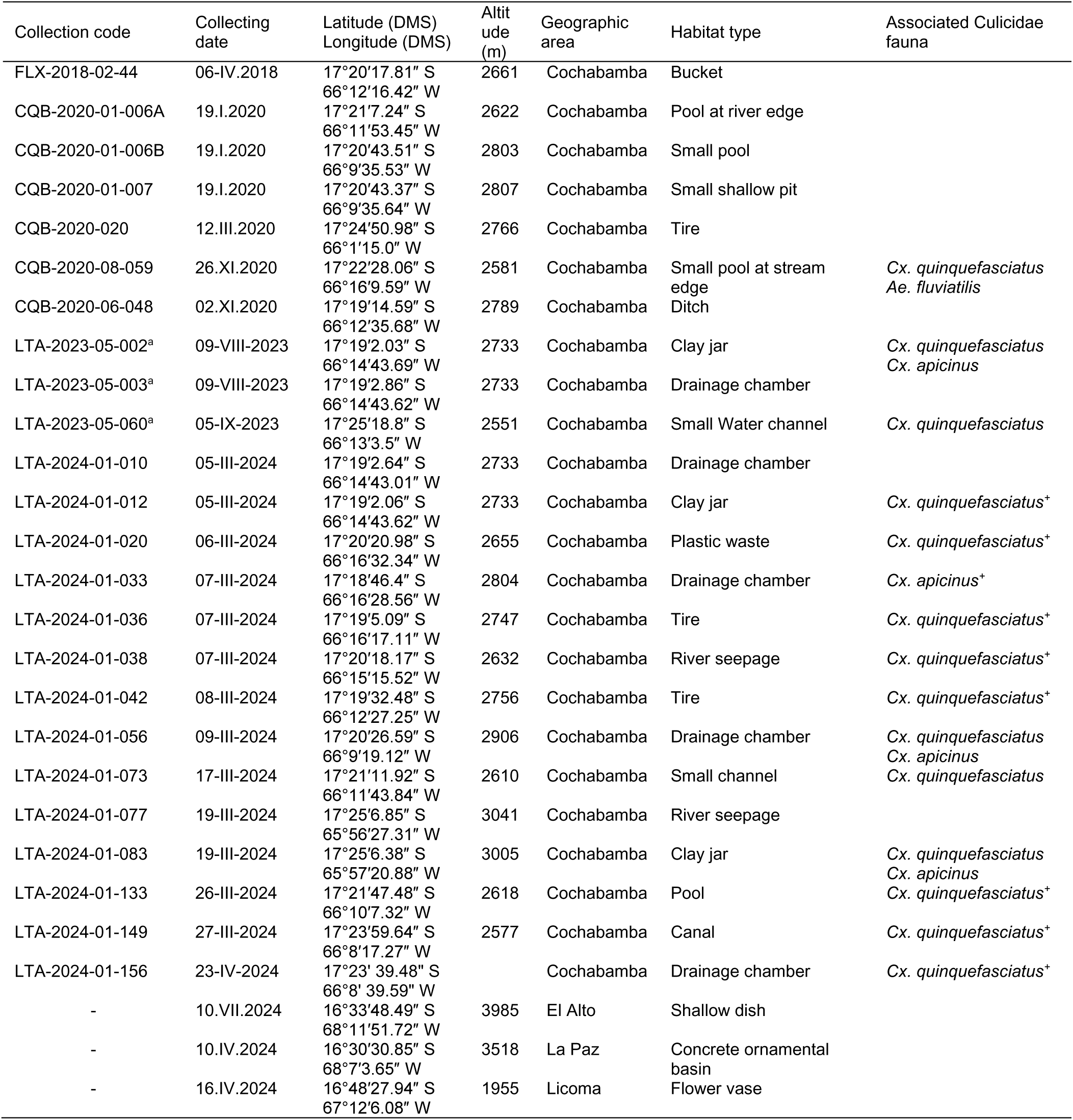
Summary of positive *Cx. acharistus* larval collections in the present study: Collection codes, dates, coordinates, habitat types, and associated Culicidae fauna. The collection code refers to the identifier used in the authors’ laboratory records at LEMUMSS (University Mayor de San Simón, Cochabamba, Bolivia). The superscript ‘a’ indicates that the sample was collected during the dry season. The sign ^+^ indicates the dominant species in the sample.

Mosquito larvae were sampled using the dipping technique (O’Malley 1995), and by direct collection from the habitat when the latter was too small to be sampled with a dip. Samples were georeferenced, and basic ecological data were recorded and stored in the VECTOBOL database (Lardeux F. *et al*. 2024a). The larvae collected were preserved in 70% alcohol until further processing for morphological identification or molecular analysis.

### Morphological identification

Fourth-instar mosquito larvae were clarified using 5% KOH, then rinsed in Marc-André solution, dehydrated through a graded ethanol series (70 −100%), immersed in clove oil, and finally mounted in Euparal (Karl Roth, Germany). Morphological identification was performed using standard dichotomous keys (Darsie 1985; Rossi Gustavo C. *et al*. 2002) and species description (Bachmann & Casal 1962; Bram 1967; Laurito *et al*. 2009). Particular attention was given to the following larval characteristics in order to distinguish *Culex (Culex) acharistus* from other members of the subgenus *Culex*. The fourth-instar larva is readily identified by a combination of distinctive morphological traits. The mentum bears approximately 25 very long, parallel-sided teeth, and the antennal setae 1-A is inserted near the middle of the subcylindrical antenna, whose base is spiculose. The abdominal integument is glabrous and includes a sclerotized plate with minute denticulations, while the siphon has a characteristic shape with a siphonal index of about 3.5, bearing four pairs of lateral setae. The pecten, located on the basal third of the siphon, comprises around 14 teeth, each with several ventral barbs. The comb consists of numerous apically fringed scales in a broad patch, and the anal segment is completely ringed by the saddle. According to Bram (1967) and Laurito *et al*. (2009), this suite of characters—the shape of the mentum, the mid-length insertion of seta 1-A, the minutely denticulate sclerotized abdominal plate, and the siphon morphology—allows for the reliable separation of *Cx. acharistus* from other *Culex (Culex)* species (FIG. 2)

**FIG. 2.**
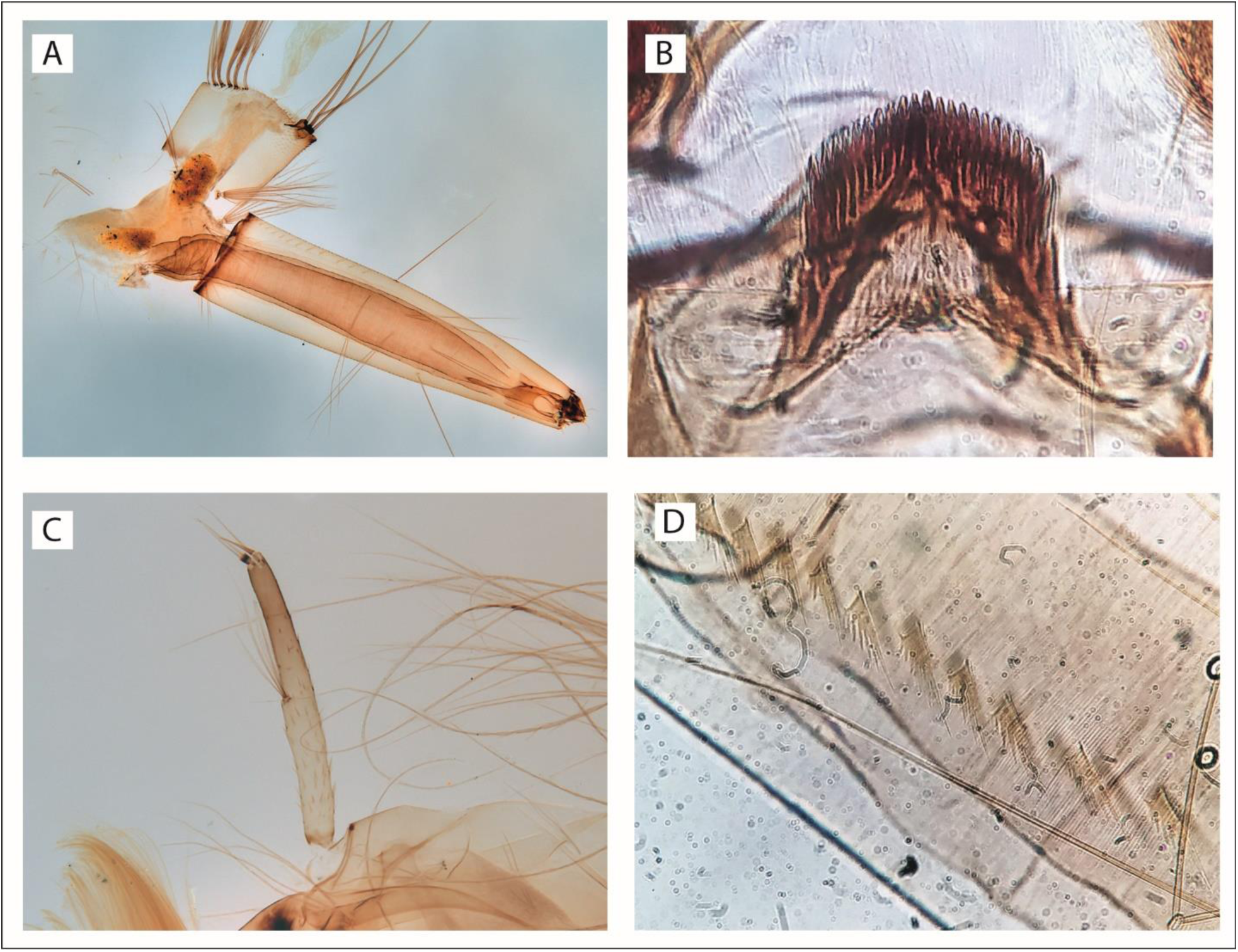
*Culex acharistus* from Bolivia. **A**, Larval siphon showing the four pairs of setae and anal segment with complete saddle (collecting site CQB-2020-01-006A in Cochabamba); **B**, Mentum (collecting site: La Paz city); **C**, Larval antenna showing setae 1-A inserted near the middle of the subcylindrical antenna, whose base is spiculose (collecting site CQB-2020-01-006A in Cochabamba); **D**, Details of pecten teeth on larval siphon (collecting site: La Paz city).

### Molecular analysis

#### DNA Extraction, PCR Amplification, and Sequencing

A total of six mosquito larvae collected in Bolivia were subjected to molecular analysis: two from El Alto (zona El Kenko; CAL01 and CAL02, as referenced in the text and FIG. 4), two from La Paz (CAL03 and CAL04), one from Licoma (CAL05), and one from Cochabamba (CQB06, collected under code CQB-2020-01-006A) (Table 1). DNA was extracted following a protocol based on CTAB/chloroform/isopropanol chemistry (Berger *et al*. 2022). The mitochondrial cytochrome c oxidase subunit I (COI) genes was amplified using primers described in Folmer *et al*. (1994): LCO1490_F (5′-GGTCAACAAATCATAAAGATATTGG-3′) and HCO2198_R (5′-TAAACTTCAGGGTGACCAAAAAATCA-3′).

Polymerase chain reactions (PCRs) were performed following protocols of Berger *et al*. (2022) and Lardeux F. *et al*. (2024b) in a total volume of 25 µL, containing 2 µL of genomic DNA (20 ng/µL), 1× Hot FIREPol® Blend Master Mix buffer, and 5 pmol of each primer. PCR amplifications were carried out using a Vapo Protect Thermocycler® under the following conditions: an initial denaturation at 95°C for 15 min; 35 cycles of 15 s denaturation at 94°C, 30 s annealing at 50°C, and 1 min extension at 72°C; followed by a final extension at 72°C for 5 min.

PCR products were purified and sequenced by Azenta Life Sciences using standard Sanger sequencing protocols.

#### Sequence editing and alignment, BLAST analysis

Forward and reverse chromatograms for each of the six individual mosquitoes were obtained as .ab1 files and visualized in MEGA 7 (Kumar *et al*. 2016). Forward and reverse sequences were compared and manually corrected by examining chromatogram peak quality, resolving ambiguities. Sequence editing and trimming were performed manually, and multiple sequence alignment was conducted using the MUSCLE algorithm implemented in MEGA 7. The final edited sequences for all six individuals were identical, resulting in a consensus haplotype of 612 base pairs (bp).

*In silico* translation of the resulting haplotype into amino acids using the invertebrate mitochondrial genetic code in MEGA 7 confirmed the presence of an intact open reading frame, consistent with a functional COI gene. BLAST analysis of the nucleotide sequence against the NCBI database returned the top four matches corresponding to *Cx. acharistus* sequences (100% query coverage, >99% sequence identity, and an e-value of 0), supporting its preliminary identification as this species. A phylogenetic analysis was subsequently conducted to confirm this assignment.

#### Sequence dataset preparation for phylogeny

In order to place the Bolivian specimens into a phylogenetic context, homologous COI sequences were retrieved from GenBank. Sequences were selected based on having a 100% query cover and greater than 98% sequence identity to the Bolivian haplotype. Forty-one sequences were initially retained. Sequences with uncertain taxonomic identification or poor annotation were excluded from the dataset. Excluded sequences were: KF919249.1, EF645645.1, MT999166.1, and KR656636.1. According to Bram (1967), *Cx. acharistus*, *Culex restuans* Theobald, 1901, *Culex brethesi* Dyar, 1919, and *Culex laticlasper* Galindo & Blanton, 1954 form a complex within the *Culex* subgenus. BLAST analysis of our six sequences indicated close matches with *Cx. acharistus* and *Cx. restuans*, two species within this complex. To further refine the phylogenetic analysis, additional COI sequences of *Cx. brethesi*—another member of this complex—were incorporated. However, only three COI sequences of *Cx. brethesi* were available in GenBank, and no sequences were found for *Cx. laticlasper.* The final dataset comprised 36 sequences, including the six Bolivian individuals. All sequences were aligned using MUSCLE and trimmed to match the 612 bp length of the Bolivian haplotype. Unique haplotypes were identified using VSEARCH v2.30.0 (Rognes *et al*. 2016) with a clustering threshold of 100% identity. The analysis identified 19 distinct haplotypes, (11 *Cx. restuans*, 5 *Cx. acharistus*, 2 *Cx. brethesi*, and 1 Bolivian haplotype constituted by the 6 individuals of the study). Although forming one haplotype, the Bolivian individuals were maintained as separate identities (CAL01-CAL05, CQB06) for clarity in the phylogenetic tree visualization.

#### Phylogenetic analysis

Phylogenetic relationships among haplotypes were reconstructed using the Maximum Likelihood (ML) method as implemented in MEGA 7. The best-fit nucleotide substitution model was determined using the “Find Best DNA/Protein Models (ML)” option of MEGA 7, and the Tamura 3-parameter model with Gamma-distributed rate variation among sites (T92+G) (Tamura 1992) was selected. ML tree inference employed an initial tree generated automatically by applying Neighbor-Joining and BioNJ algorithms to a matrix of pairwise distances estimated by the Maximum Composite Likelihood (MCL) method. Branch support was evaluated through 500 bootstrap replicates. The Maximum Likelihood (ML) final tree was built with a final alignment of 611 bp. Evolutionary analyses were conducted in MEGA 7.

#### Acronyms of repositories

LEMUMSS = Laboratorio de Entomología Medica de la Universidad Mayor de San Simón, Cochabamba, Bolivia.

INLASA = Laboratorio de Entomología Medica del Instituto Nacional de Laboratorios de Salud, La Paz, Bolivia.

#### Cited material

BOLIVIA – **Cochabamba department •** Tiquipaya city; 17°20′17.81″ S; 66°12′16.42″ W; 2661 m; 06 Apr. 2018; F. Lardeux leg.; LEMUMSS, FLX-2018-02-44 – **Cochabamba department •** 17°21′7.24″ S; 66°11′53.45″ W; 2622 m; 19 Jan. 2020; C. Quezada leg.; LEMUMSS, CQB-2020-01-006 – **Cochabamba department •** 17°20′43.51″ S; 66°9′35.53″ W; 2803 m; 19 Jan. 2020; C. Quezada leg.; LEMUMSS, CQB-2020-01-006B – **Cochabamba department •** 17°20′43.37″ S; 66°9′35.64″ W; 2807 m; 19 Jan. 2020; C. Quezada leg.; LEMUMSS, CQB-2020-01-007 – **Cochabamba department •** 17°24′50.98″ S; 66°1′15.0″ W; 2766 m; 12 Mar. 2020; C. Quezada leg.; LEMUMSS, CQB-2020-020 – **Cochabamba department •** 17°22′28.06″ S; 66°16′9.59″ W; 2581 m; 26 Nov. 2020; C. Quezada leg.; LEMUMSS, CQB-2020-08-059 – **Cochabamba department •** 17°19′14.59″ S; 66°12′35.68″ W; 2789 m; 02 Nov. 2020; C. Quezada leg.; LEMUMSS, CQB-2020-06-048 – **Cochabamba department •** 17°19′2.03″ S; 66°14′43.69″ W; 2733 m; 09 Aug. 2023; L. Torrez leg.; LEMUMSS, LTA-2023-05-002 – **Cochabamba department •** 17°19′2.86″ S; 66°14′43.62″ W; 2733 m; 09 Aug. 2023; L. Torrez leg.; LEMUMSS, LTA-2023-05-003 – **Cochabamba department •** 17°25′18.8″ S; 66°13′3.5″ W; 2551 m; 05 Nov. 2023; L. Torrez leg.; LEMUMSS, LTA-2023-05-060 – **Cochabamba department •** 17°19′2.64″ S; 66°14′43.01″ W; 2733 m; 05 Mar. 2024; L. Torrez leg.; LEMUMSS, LTA-2024-01-010 – **Cochabamba department •** 17°19′2.06″ S; 66°14′43.62″ W; 2733 m; 05 Mar. 2024; L. Torrez leg.; LEMUMSS, LTA-2024-01-012 – **Cochabamba department •** 17°20′20.98″ S; 66°16′32.34″ W; 2655 m; 06 Mar. 2024; L. Torrez leg.; LEMUMSS, LTA-2024-01-020 – **Cochabamba department •** 17°18′46.4″ S; 66°16′28.56″ W; 2804 m; 07 Mar. 2024; L. Torrez leg.; LEMUMSS, LTA-2024-01-033 – **Cochabamba department •** 17°19′5.09″ S; 66°16′17.11″ W; 2747 m; 07 Mar. 2024; L. Torrez leg.; LEMUMSS, LTA-2024-01-036 – **Cochabamba department •** 17°20′18.17″ S; 66°15′15.52″ W; 2632 m; 07 Mar. 2024; L. Torrez leg.; LEMUMSS, LTA-2024-01-038 – **Cochabamba department •** 17°19′32.48″ S; 66°12′27.25″ W; 2756 m; 08 Mar. 2024; L. Torrez leg.; LEMUMSS, LTA-2024-01-042 – **Cochabamba department •** 17°20′26.59″ S; 66°9′19.12″ W; 2906 m; 09 Mar. 2024; L. Torrez leg.; LEMUMSS, LTA-2024-01-056 – **Cochabamba department •** 17°21′11.92″ S; 66°11′43.84″ W; 2610 m; 17 Mar. 2024; L. Torrez leg.; LEMUMSS, LTA-2024-01-073 – **Cochabamba department •** 17°25′6.85″ S; 65°56′27.31″ W; 3041 m; 19 Mar. 2024; L. Torrez leg.; LEMUMSS, LTA-2024-01-077 – **Cochabamba department •** 17°25′6.38″ S; 65°57′20.88″ W; 3005 m; 19 Mar. 2024; L. Torrez leg.; LEMUMSS, LTA-2024-01-083 – **Cochabamba department •** 17°21′47.48″ S; 66°10′7.32″ W; 2618 m; 26 Mar. 2024; L. Torrez leg.; LEMUMSS, LTA-2024-01-133 – **Cochabamba department •** 17°23′59.64″ S; 66°8′17.27″ W; 2577 m; 27 Mar. 2024; L. Torrez leg.; LEMUMSS, LTA-2024-01-149 – **Cochabamba department •** 17°23’ 39.48” S; 66°8’ 39.59” W; 23 Apr. 2024; L. Torrez leg.; LEMUMSS, LTA-2024-01-156 – **La Paz department •** El Alto city; 16°33′48.49″ S; 68°11′51.72″ W; 3985 m; 10 jul. 2024; P. Oporto leg.; INLASA – **La Paz department •** La Paz city; 16°30′30.85″ S; 68°7′3.65″ W; 3518 m; 10 Apr. 2024; C. Aliaga leg.; INLASA – **La Paz department • Licoma;** 16°48′27.94″ S; 67°12′6.08″ W; 1955 m; J. Barrera leg.; 16 Apr. 2024; INLASA.

## Results

### Field samples

*Culex acharistus* has been collected in Bolivia from a variety of larval habitats, including peri-domestic containers as well as natural and artificial water bodies (Table 1, FIG. 3).

**FIG. 3.**
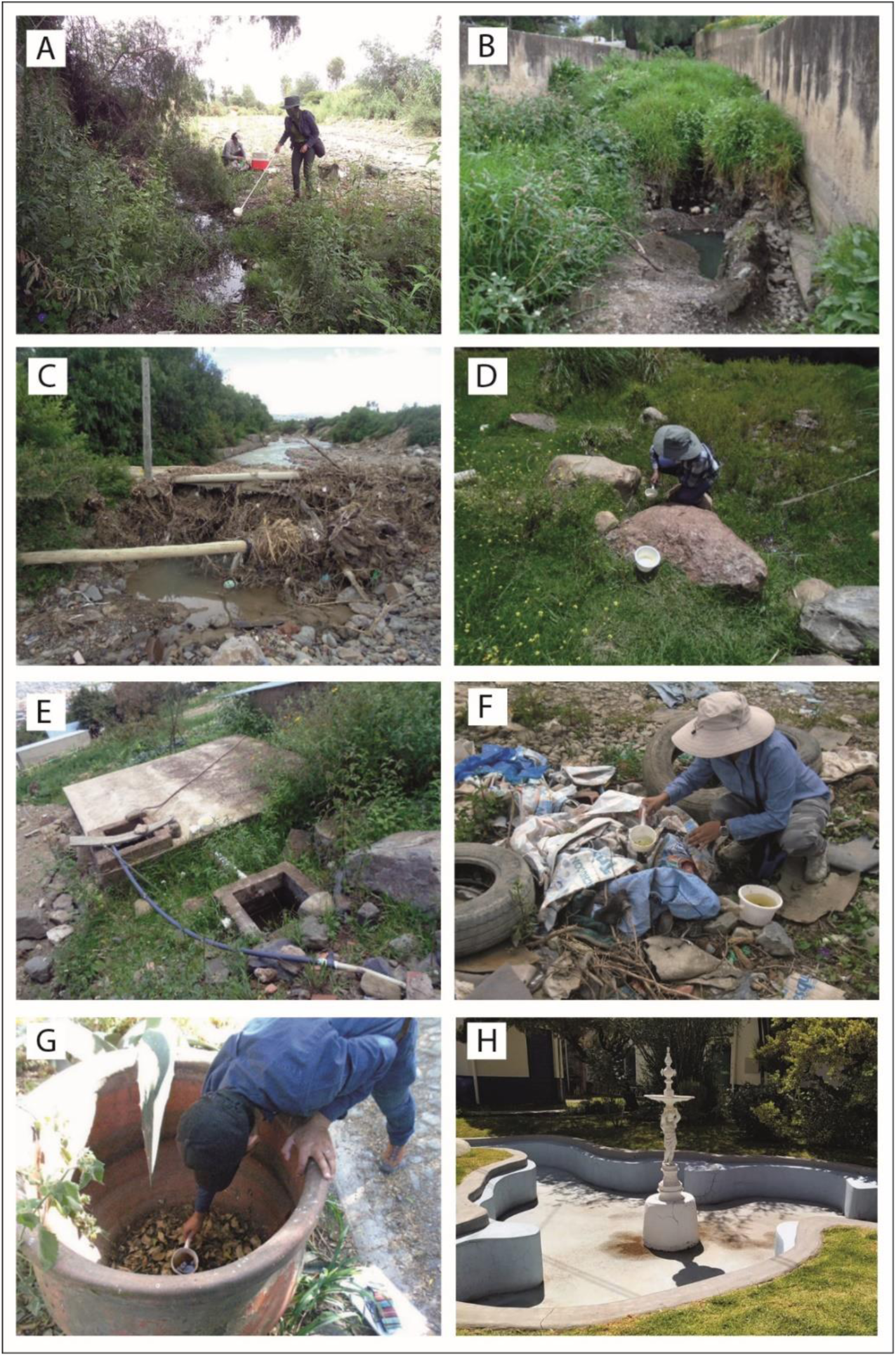
Some Bolivian larval habitats of *Cx. acharistus*. **A**, Small pool at stream edge (CQB-2020-08-059); **B**, Pool (LTA-2024-01-133); **C,** River seepage (LTA-2024-01-038); **D**, River seepage (LTA-2024-01-077); **E**, Drainage chamber (LTA-2024-01-056); **F**, Plastic waste (LTA-2024-01-020); **G**, Clay jar (LTA-2023-05-002); **H,** Concrete ornamental basin. The codes correspond to those listed in Table 1. All people shown in the photographs are some of the article’s authors.

In the La Paz/El Alto region, the first collection site was a small, disused, circular concrete ornamental basin (≈20 m²) located within the grounds of the Instituto Nacional de Laboratorios de Salud (INLASA) in central La Paz (FIG. 3, G). The second site, in the El Kenko district of El Alto, was a shallow dish used as a water source for a dog. In Licoma, specimens were collected from a flower vase placed in the municipal cemetery. In these three occasions, *Cx. acharistus* was collected alone.

In the Cochabamba area, during the three surveys conducted in 2020, 2023, and 2024, a total of 166 larval habitats were sampled, of which 24 were colonized by *Cx. acharistus* (Table 1). Three other mosquito species were also identified in these habitats: *Aedes fluviatilis* (Lutz, 1904)*, Culex apicinus* Philippi, 1865, and *Culex quinquefasciatus* Say, 1823. Across all samples, *Cx. acharistus* was found alone on 12 occasions; in association with *Cx. quinquefasciatus* on 10 occasions; with *Cx. apicinus* on one occasion; with both *Cx. quinquefasciatus* and *Cx. apicinus* on three occasions; and with both *Cx. quinquefasciatus* and *Ae. fluviatilis* on one occasion. (Table 1). Overall, when *Cx. acharistus* co-occurs with *Cx. quinquefasciatus*, the latter tends to be the dominant species within the sample. *Culex acharistus* was collected in the Cochabamba area during both the rainy and dry seasons, although it was less frequently encountered in the dry season (Table 1).

### Phylogenic analysis

The six *Cx. acharistus* COI sequences from Bolivia were deposited in GenBank. The sequences from specimens CAL01 and CAL02 from El Alto city correspond to accession numbers PX131006-1 and PX132435-1; CAL03 and CAL04 from La Paz city correspond to PX132436-1 and PX132434-1; CAL05 from Licoma corresponds to PX132438-1; and CQB06 from Cochabamba city corresponds to PX132437-1. Phylogenetic analysis of the COI gene sequences revealed a single haplotype among all six Bolivian specimens, identified as *Cx. acharistus* based on both BLAST results and phylogenetic clustering. The analysis included 24 sequences, comprising haplotype sequences from *Cx. acharistus*, *Cx. restuans*, *Cx. brethesi*, and the 6 references species. In the ML tree the Bolivian sequences consistently grouped within the *Cx. acharistus* clade, showing high bootstrap support (85%) for the placement of these sequences (FIG. 4). The tree structure also revealed a clear separation between *Cx. acharistus* and *Cx. restuans*, with the latter forming a distinct clade supported by moderate bootstrap values (72%). Additionally, *Cx. brethesi* was well supported as a separate group (bootstrap 98%), distinct from both *Cx. acharistus* and *Cx. restuans*. These results confirm the identification of the Bolivian specimens as *Cx. acharistus* and provide further insight into the phylogenetic relationships within this subgroup of the *Culex* genus.

**FIG. 4.**
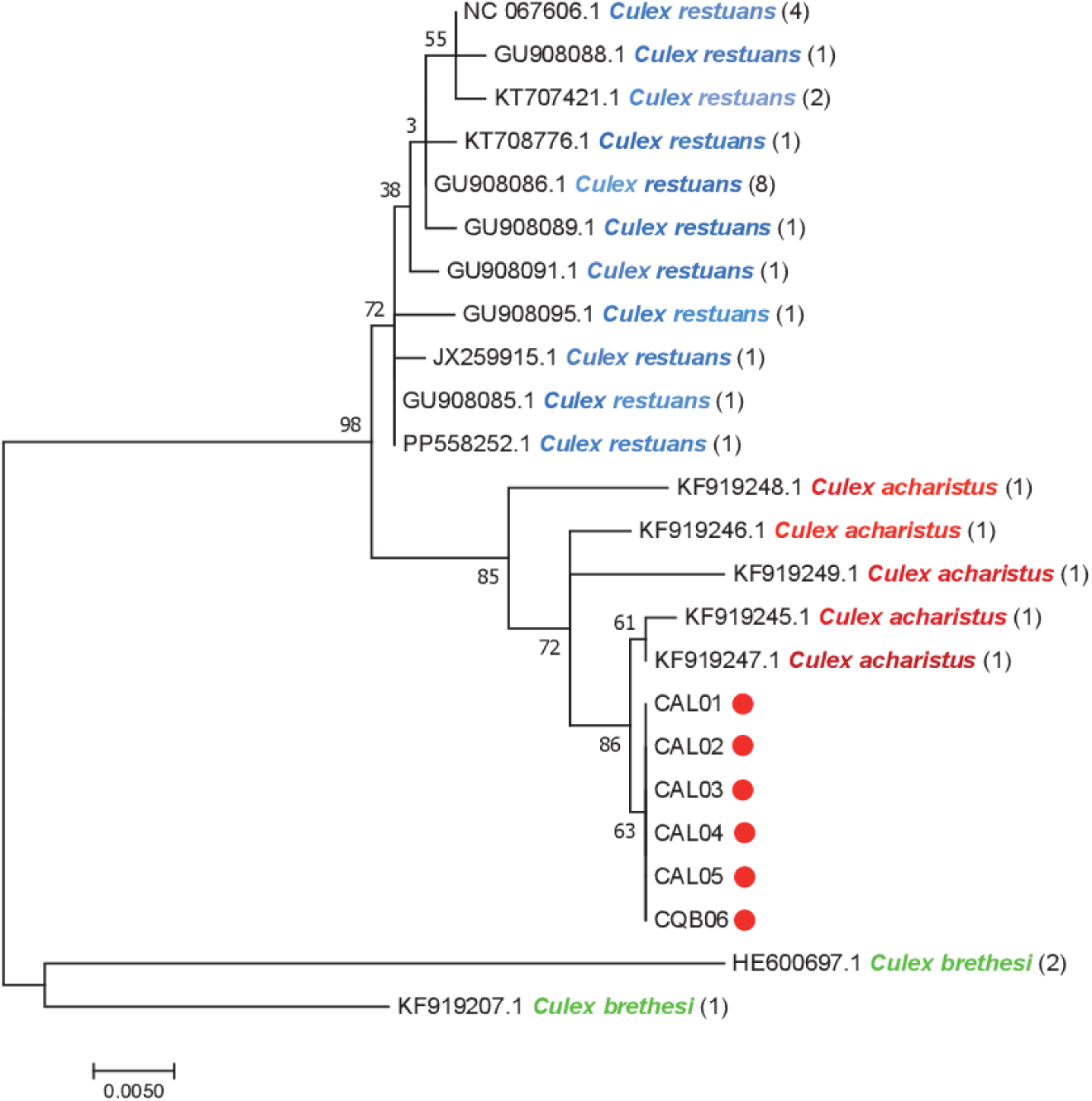
Maximum Likelihood tree. The tree with the highest log likelihood (−1253.38) is shown. Numbers in brackets indicate the number of haplotypes. Bolivian specimens are represented by red dots. All *Cx. acharistus* and *Cx. brethesi* specimens are from Argentina, and all *Cx. restuans* specimens are from North America (USA, Canada).

The percentage of trees in which the associated taxa clustered together is shown next to the branches. Initial tree(s) for the heuristic search were obtained automatically by applying Neighbor-Join and BioNJ algorithms to a matrix of pairwise distances estimated using the Maximum Composite Likelihood (MCL) approach, and then selecting the topology with superior log likelihood value. A discrete Gamma distribution was used to model evolutionary rate differences among sites (5 categories (+G, parameter = 0.0500)). The tree is drawn to scale, with branch lengths measured in the number of substitutions per site. The analysis involved 24 nucleotide sequences. Codon positions included were 1st+2nd+3rd+Noncoding. All positions containing gaps and missing data were eliminated. There was a total of 611 positions in the final dataset tree based on an alignment of 611 nucleotides of the COI mitochondrial gene. Evolutionary analyses were conducted in MEGA7

## Discussion

*Culex (Culex) acharistus* was first described by Root in 1927 from specimens collected in March 1925 from Agua Limpa, Brazil (Root 1927). The description was based on female, male and male genitalia, but the immature stages were not preserved and no holotype was designated. Subsequent contributions progressively clarified the species’ morphology: Lane (1953) provided a partial description of the pupa; Bachmann & Casal (1962) gave the first descriptions of the larva and pupa from Argentina, albeit briefly and with incomplete illustrations; Forattini & Rabello (1965) later offered a more comprehensive characterization of the pupal stage, and Bram (1967) refined the species’ diagnosis allowing clearer separation from morphologically similar species. A male lectotype was later designated by Stone & Knight (1957) and more recently, Laurito *et al*. (2009) re-described the immatures and adult stages of the species. Based on taxonomy literature, the Bolivian specimens were unambiguously assigned to *Cx. acharistus*.

The molecular analysis based on the mitochondrial COI gene supports this identification. Al Bolivian sequences clustered with *Cx. acharistus* from northeastern Argentina (KF919245–KF919249) with strong bootstrap support. Moreover, COI barcoding proved effective for distinguishing closely related *Culex* species, as shown by the clear separation of *Cx. acharistus* from taxa such as *Cx. restuans* and *Cx. brethesi* in the ML tree, consistent with the *Cx. acharistus* species’ validity as recognized by Laurito *et al*. (2013). The absence of sequence divergence among the six Bolivian individuals suggests limited mitochondrial variability in the sampled population, potentially reflecting a low local genetic diversity. Moreover, the close genetic similarity between the Bolivian and Argentinian specimens suggests a low intraspecific divergence within *Cx. acharistus* across the southern Neotropical region. No COI sequences of *Cx. acharistus* from other neighboring countries where the species is known to occur were available for direct comparison.

The detection of *Cx. acharistus* in Bolivia increases the number of species recorded in the *Culex* subgenus in the country to at least 17, based on current bibliographic records and the examination of voucher specimens deposited in the entomological collections of INLASA (La Paz) and LEMUMSS (Cochabamba) (VECTOBOL 2025). However, this updated figure should be interpreted with caution, as species-level identifications within the subgenus *Culex* are often hindered by morphological similarities, taxonomic ambiguities, and the lack of recent comprehensive revisions.

In spite of increased taxonomic efforts, *Cx. acharistus* remains one of the Neotropical *Culex* species that lack comprehensive study. This knowledge gap is particularly relevant given the potential medical importance of the species. During an Eastern Equine Encephalitis epizootic in horses in Chaco Province, Argentina, in 1988, *Cx. acharistus* was the most abundant mosquito species observed, raising suspicion about its role in virus transmission (Avilés *et al*. 1989). More recently, it has been found naturally infected with Bunyamwera virus in Formosa Province, Argentina (Gallardo *et al*. 2019), suggesting a possible but as yet unconfirmed involvement in arbovirus transmission cycles.

Ecologically, *Cx. acharistus* is restricted to South America, where it is widely distributed, with confirmed occurrences in Brazil (Root 1927; Navarro da Silva & Leuch 1996), Chile (Bram 1967), Peru (Ayala-Sulca *et al*. 2025)Argentina ((Almirón W. R. *et al*. 1995; Campos R. E. & Marcia 1998; Rossi G. C. 2000; Cardo *et al*. 2024), Colombia (Bram 1967; Rosero-Garcia *et al*. 2017) Uruguay (Rossi G. C. 2014; Campos R. *et al*. 2025) and, as reported here for the first time, Bolivia.

The species is primarily associated with mid- to high-elevation environments (1200 - 2600 m.a.s.l.) (Rosero-Garcia *et al*. 2017; Ayala-Sulca *et al*. 2025), but it has also been reported at lower altitudes, including a few hundred meters and down to sea level in Chile (Bram 1967; Angulo & Olivares 1993), Peru (Ayala-Sulca *et al*. 2025) and Brazil (Navarro da Silva & Leuch 1996; Barbosa *et al*. 2003), and temperate to cool subtropical climates (Grech *et al*. 2023). Its occurrence both in the Andes (Rosero-Garcia *et al*. 2017) and Patagonia (Almirón W. R. *et al*. 1995; Rossi G. C. & Vezzani 2011; Grech *et al*. 2019) demonstrates its adaptation to colder environments. Its presence in the Bolivian Andes is therefore consistent with its known altitudinal and climatic distribution elsewhere in the region, and represents for the El Alto record at ≍4000 m, one of the highest elevation records reported for the species.

Larval ecology studies indicate that *Cx. acharistus* displays moderate ecological plasticity. Immature stages occur in a wide range of natural aquatic habitats, including marshy floodplains, stream margins, rain-filled rock pools, animal footprints and temporary or permanent ground pools with or without aquatic vegetation (Laurito *et al*. 2009; Linares *et al*. 2016). These habitats can be located in both shaded and sunlit environments. The species also colonizes artificial containers, such as discarded tires, cisterns, swimming pools, plastic or metal receptacles, and ditches—especially in peri-domestic or urban settings (Rossi G. C. & Almirón 2004; Achaga & Vezzani 2024; Cardo *et al*. 2024), suggesting increasing adaptation to anthropogenic environments. In Bolivia, specimens were collected from both natural and artificial containers, in agreement with patterns observed elsewhere in its distribution range (Grech *et al*. 2023).

In Argentina, *Cx. acharistus* larvae frequently occur in sympatry with other *Culex* species, particularly *Cx. quinquefasciatus*, *Cx. bidens* Dyar, 1922, *Cx. dolosus* (Lynch Arribálzaga, 1891), *Cx. maxi* Dyar, 1928, and *Cx. apicinus* (Almirón W. R. & Brewer 1994; Almirón W. M. & Brewer 1996; Díaz Nieto *et al*. 2020). In Bolivia, it was also found in association with *Cx. quinquefasciatus*, *Cx. apicinus*, and *Ae. fluviatilis*, mirroring patterns observed in Argentina. These associations suggest not only shared ecological preferences but also potential niche partitioning among species.

In Bolivia, *Cx. acharistus* may be more widespread than previously recognized. Our field observations in Cochabamba indicate that once targeted searches are conducted, the species can be readily detected, particularly in urban and peri-urban settings. This underscores the likelihood that *Cx. acharistus* is under-reported at the country level, due to limited sampling rather than true rarity.

This study extends the known distribution of genetically confirmed *Cx. acharistus* to Bolivia and contributes to its molecular characterization. This is particularly relevant given the taxonomic challenges and complexity within the *Culex* genus, where cryptic species and morphological convergence often complicate species-level identification. The congruence between morphological and molecular data in this study reinforces the reliability of both approaches and highlights the importance of integrative taxonomy in entomological biodiversity assessments.

The confirmed presence of *Cx. acharistus* in Bolivia adds valuable information to the relatively poorly documented mosquito fauna of the country, particularly in highland regions. This record not only suggests the possible presence of other, yet undetected, highland *Culex* species but also underscores the need for continued, targeted entomological surveys. Furthermore, these findings provide a foundation for future ecological and epidemiological studies, particularly in the context of climate change, which may alter mosquito distributions and associated vector-borne disease risks.

## Acknowledgements

We would like to thank Dra. P. Oporto from the Entomology and Parasitology Laboratory at INLASA, La Paz for providing the mosquito specimens collected in El Kenko (El Alto), and Technician J. Barrera from SEDES-La Paz for those collected in Licoma. We thank S. Quispe and C. Zubieta, from the Entomology Laboratory LEMUMSS of Cochabamba who helped in collecting the mosquitoes in the Cochabamba area. We also thank C. Barnabé (UMR INTERTRYP, IRD-CIRAD-UM-II) for his help in analyzing the molecular data, including sequence editing, phylogenetic tree construction, and interpretation.

